# Capturing individual variation in children’s oscillatory brain activity during nREM sleep

**DOI:** 10.1101/2025.05.02.651817

**Authors:** Verna Heikkinen, Susanne Merz, Riitta Salmelin, Sampsa Vanhatalo, Leena Lauronen, Mia Liljeström, Hanna Renvall

## Abstract

Human brain dynamics are highly unique between individuals: functional neuroimaging studies have recently described functional features that can be used as neural fingerprints. As such studies have generally been performed on adults, it is unclear if the results are applicable during brain development. Here we applied Bayesian reduced-rank regression (BRRR), to extract low-dimensional representations of electroencephalography (EEG) power spectra measured from normally developing children at different ages. The clinical EEG recordings represented different sleep stages in a pediatric population (N=782) of ages varying from 6 weeks to 19 years. The representations learned within specific sleep stages successfully separated between the subjects. Furthermore, a model trained with data across sleep stages generalized to unseen data, suggesting that informative neural fingerprints could be extracted. Fingerprint stability increased with the age of the subjects. In comparison to correlation-based measures in differentiating individuals from each other, the BRRR model performed better, highlighting the usefulness of the dimensionality reduction for data obtained in demanding clinical settings. While further studies are needed to address the possible non-linear maturation effects over developmental periods, our results demonstrate, for the first time, the existence of stable within-session neurofunctional fingerprints in pediatric populations.

## Introduction

The human cerebral cortex demonstrates salient, intrinsic oscillatory activity that can be characterized non-invasively with electroencephalography (EEG) and magnetoencephalography (MEG). This spontaneous activity is altered in many neurological diseases, and it is therefore frequently studied in both research and clinical settings in pediatric and adult populations [26, 52, 65]. Despite high variability of oscillatory activity patterns between individuals [28, 33], many typical EEG or MEG features appear to be very consistent within adult subjects over time. Indeed, recent MEG and fMRI studies have identified stable neural patterns within individuals’ own data samples, referred commonly as brain fingerprints, among functional connectomes and broad-band oscillatory activation patterns [2, 11, 14, 21, 22, 27, 35, 36, 65], even generalizing across experimental tasks [21, 22, 36]. Some of the individual variation has been linked to genetic factors [3, 32, 36, 56], but as the stability of the individual features decreases with aging [14] and disease [12, 53], the individual patterns appear not exclusively determined by genetics. While the studies have revealed highly individual yet stable functional brain dynamics in healthy adults, it is not clear whether the findings generalize to pediatric populations.

Significant changes in M/EEG activity appear during brain maturation [4]. Low frequency (< 4 Hz) oscillations decrease in power with age, both during sleep [19] and resting wakefulness [10, 45, 63], while activity in higher frequency bands increases, as demonstrated by the emergence of theta (4 – 10 Hz) and sigma oscillations (12 – 15 Hz) [48] and by changes in the gamma (*>* 30 Hz) range [8]. In addition, the peak frequency in the alpha-range (8 – 12 Hz) starts to increase after around 16 months of age [8, 10]. Sleep-related M/EEG features, such as spindle density, are also associated with age, although the relationship appears non-linear [46].

While maturation-related changes in EEG patterns generally follow a predictable trajectory, the variability is high between individuals [5, 46, 57]. Interestingly, the difference between the chronological age of an individual and that predicted by a person’s structural and functional neuroimaging features, or the so-called brain age, has been connected to cognitive functions both in healthy [20] and diseased populations [51]. Neuroimaging features thus show potential as cognitive, or brain health, biomarkers. The stability of these individual features, or the lack thereof, could similarly serve as a surrogate biomarker of risk of atypical development. However, before examining clinically relevant deviations, the individual stability and variation of neuroimaging features over brain maturation in normally developing children needs to be addressed. Furthermore, any approach aimed for clinical use should be validated with real clinical data [54], and its flexibility addressed in the varying and often noisy circumstances encountered in clinical practice.

Sleep EEG offers an excellent measure for studying the effects of individual variation and age in large children cohorts. It is cheap and easy to administer even for infants and small children who otherwise may be uncooperative during wakeful recordings. It is widely administered in clinical practice, e.g., in epilepsy diagnostics in the pediatric and adult populations [26, 52], thus offering large real-world data sets. Sleep EEG is typically divided to rapid eye movement (REM) and non-REM (nREM) stages, the latter constituting three sub-stages (N1–N3) according to the depth of sleep. Akin to task-related and resting-state paradigms, the sleep EEG has been shown to be very stable in adults [13], with some trait-like features reported in adolescents and teens despite the ongoing neurodevelopmental changes [55].

Here we aimed to address whether within-session brain fingerprints, based on power spectral features in EEG recordings, can be extracted and generalized in children during maturation. For this, we applied a machine-learning based approach to analyze power spectra in clinical sleep-EEG recordings from a healthy pediatric population ranging from infants to adolescents (N = 782). Attempts to explain high-dimensional and correlated functional brain responses along with surrogate identifiers, such as age or sex, give rise to complex but sparse data models. Such problems are increasingly tackled with probabilistic learning and latent variable models [38, 66]. One variant is latent-noise Bayesian reduced-rank regression (BRRR) [24, 42], which has shown promising results in discovering fingerprint-like features in adults’ resting-state MEG data [27, 36]. The latent-noise formulation encodes the assumption that noise affects both the target and explanatory variables similarly and thus allows to exploit the correlation structures present in the data. Previous approaches to find individually stable features from functional neuroimaging data have generally relied on correlations between different measurements of the same participants. While correlation approaches can reliably identify individuals within specific subject groups, such as patients and controls [12], it has remained uncertain which factors contribute to the differentiability of any individual subject.

We sought to find the latent representation of the EEG data that would maximally differentiate subjects from each other while preserving individual stability over data samples. The formulated model was validated by using it to differentiate individuals within and between different sleep stages, as well as to classify data from unseen participants. We compared the results to those obtained using a correlation-based method [11] and examined the effect of age and sex on the differentiability of the subjects. We hypothesized that sleep patterns would become increasingly stable with age and thus allow for better individual data matching along with maturation; sleep-EEG patterns become increasingly adult-like during late childhood and adolescence [31].

We demonstrate here high individual variation and within-session differentiability of children using their sleep-EEG bandpower features. The BRRR-based low-dimensional representation resulted in individual fingerprints that were more stable than those obtained through a correlation approach. In addition, the stability of the low-dimensional fingerprint was dependent on the age, and to a lesser degree, sex of the participants.

## Results

### Spectral power patterns change non-linearly during childhood

The dataset consisted of 19-channel clinical EEG recordings of non-REM sleep in 782 healthy children aged between 6 weeks and 19 years (mean age ± SD: 4.6 ± 4.3 years). The analyzed part of the recording encompassed 900 seconds of spontaneous sleep data. The first third (300 s, categorized as N1) consisted of resting EEG before the onset of N2 sleep, followed by 600 s of EEG after the first signs of N2 sleep. From the EEG recordings, we calculated power spectral density estimates (PSDs) for six data segments: two 60s segments that were categorized as N1 sleep, and four 60s segments that were categorized as N2 sleep.

Age-related differences were observed in the PSD patterns of the N1 and N2 sleep segments (Figure 1). For both sleep stages, the overall spectral power, as indicated by the area under the curve (AUC) of the power spectrum, decreased with age (*F* = 114.6, *p <* 0.0001). This decline was not linear from infancy to adolescence: the overall power across frequencies increased during the first year of life (*p <* 0.0001), possibly related to skull ossification, and started to decrease from 1.3 – 2 years onwards (Figure 1B). Both the youngest (ages 6 weeks - 6 months) and the oldest age groups (11.4 – 19 yrs) differed significantly from the other age groups (*p <* 0.0001).

**Figure 1.**
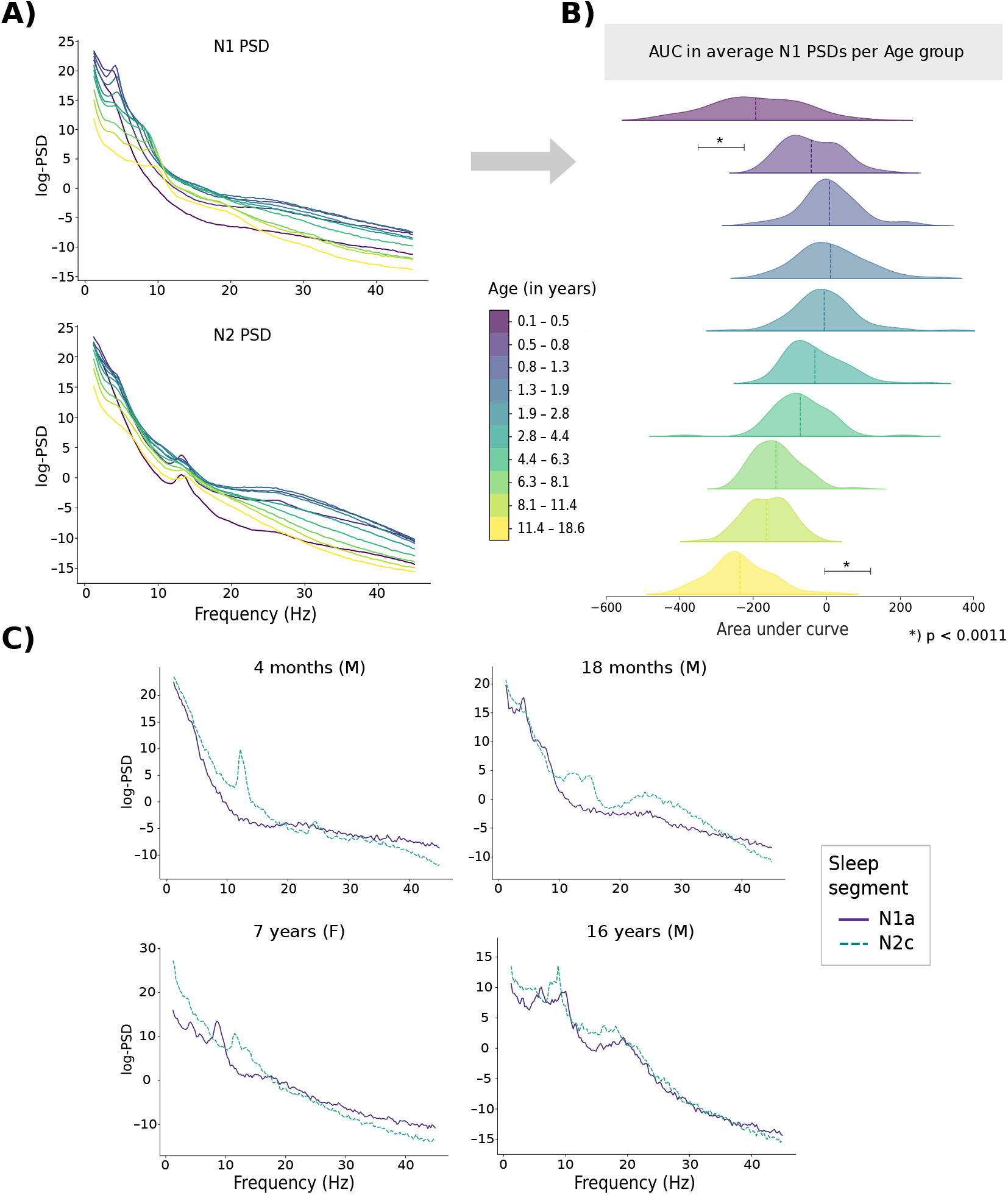
**A)** Grand average PSD estimates based on 1-minute segments of N1 and N2 sleep, calculated for ten equal-sized age groups. PSDs are represented on a logarithmic scale. **B)** The N1-PSD area under curve (AUC) demonstrates the increase in the total spectral power during the first year of life and the following decrease. The AUC values of the oldest and youngest age group differed significantly from all the other age groups; significant p-values (Bonferroni-corrected) are indicated by stars. For the rest of the groups, the difference in means was significant against the other age groups except for their immediate neighbors. **C)** PSDs averaged over all channels in four example subjects across 1-min sleep state segments (one from N1 sleep stage, one from N2 sleep stage). F= female, M = male. N1a and N2c refer to the first and third segments within the respective sleep stages.

Within the N1 sleep stage, there were systematic differences between different data segments in 6/10 of the age groups (*p <* 0.01), especially at frequencies above 26 Hz. Within the N2 sleep stage, differences also appeared in 6/10 age groups (*p <* 0.01), showing trend-like increases or decreases as the recording progressed in time (Figure 2). Overall, the spectral patterns of sleep varied across different age groups (Figure 1A) and participants (Figure 1C). In young infants (*<* 3 months), the PSDs during both sleep stages resembled non-periodic 1/f activity. Oscillatory phenomena emerged with maturation: the N1 and N2 segments within individuals differed from each other due to, e.g., emergence of sleep spindles, beta activity, and presence of low-frequency oscillatory activity (Figure 1C). In all age groups, the sleep stages differed from each other at around 12 – 15 Hz, i.e., the sleep spindle frequency range (*p <* 0.01). Despite the prominent age-related differences in the sleep PSDs, these salient global differences were not always present at the individual level (Figure 1C).

**Figure 2.**
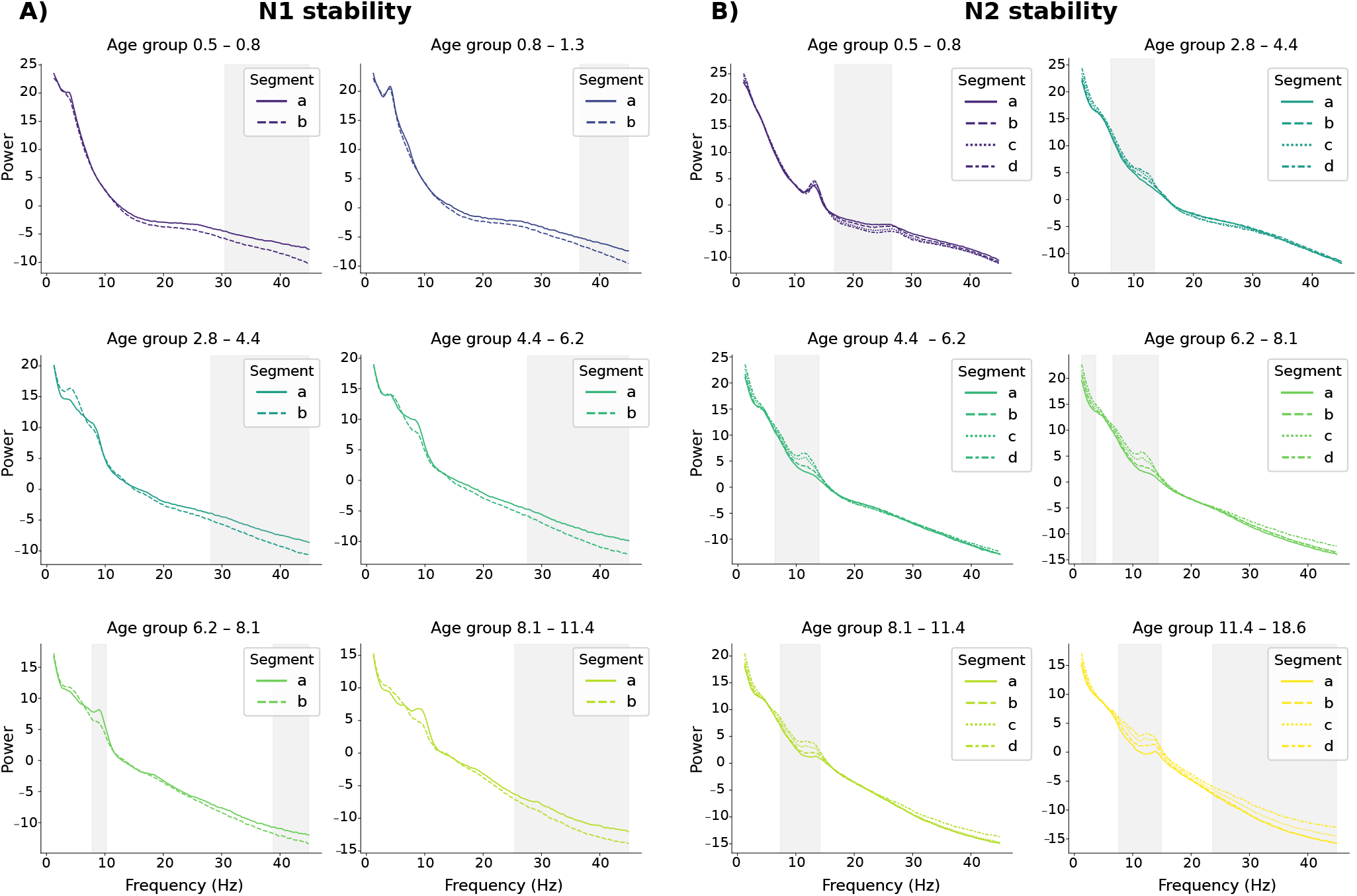
Stability of the PSD data segments across age groups during N1 (**A**) and N2 sleep (**B**). Note the different age groups in Figures A and B. Clusters indicating significant differences (*p <* 0.01) between segments within sleep stages are highlighted. The segments are marked chronologically (a–b, a–b–c–d).

### Cortical fingerprints in children can be identified from EEG spectral power

To identify individual brain fingerprints that would best discriminate between subjects, we applied the BRRR algorithm to features obtained from the PSDs. We first calculated the bandpower over 13 logarithmically widening frequency bands between 1 and 43 Hz (see Figure 3A–B). The relative bandpower in each of the 13 frequency bands was then used as a feature in the BRRR algorithm. Differentiation of subjects was carried out both within and between sleep stages. Due to the evident maturation effects, we applied the BRRR model first to all participants (N = 782; Group ‘All’) and then separately to participants who were older than seven years (N = 216; Group ‘*>*7-year-olds’) and compared the performance and latent space structures of the models. We hypothesized that the model trained using older participants could provide more stable fingerprints compared to the model using all participants, especially across sleep stages.

**Figure 3.**
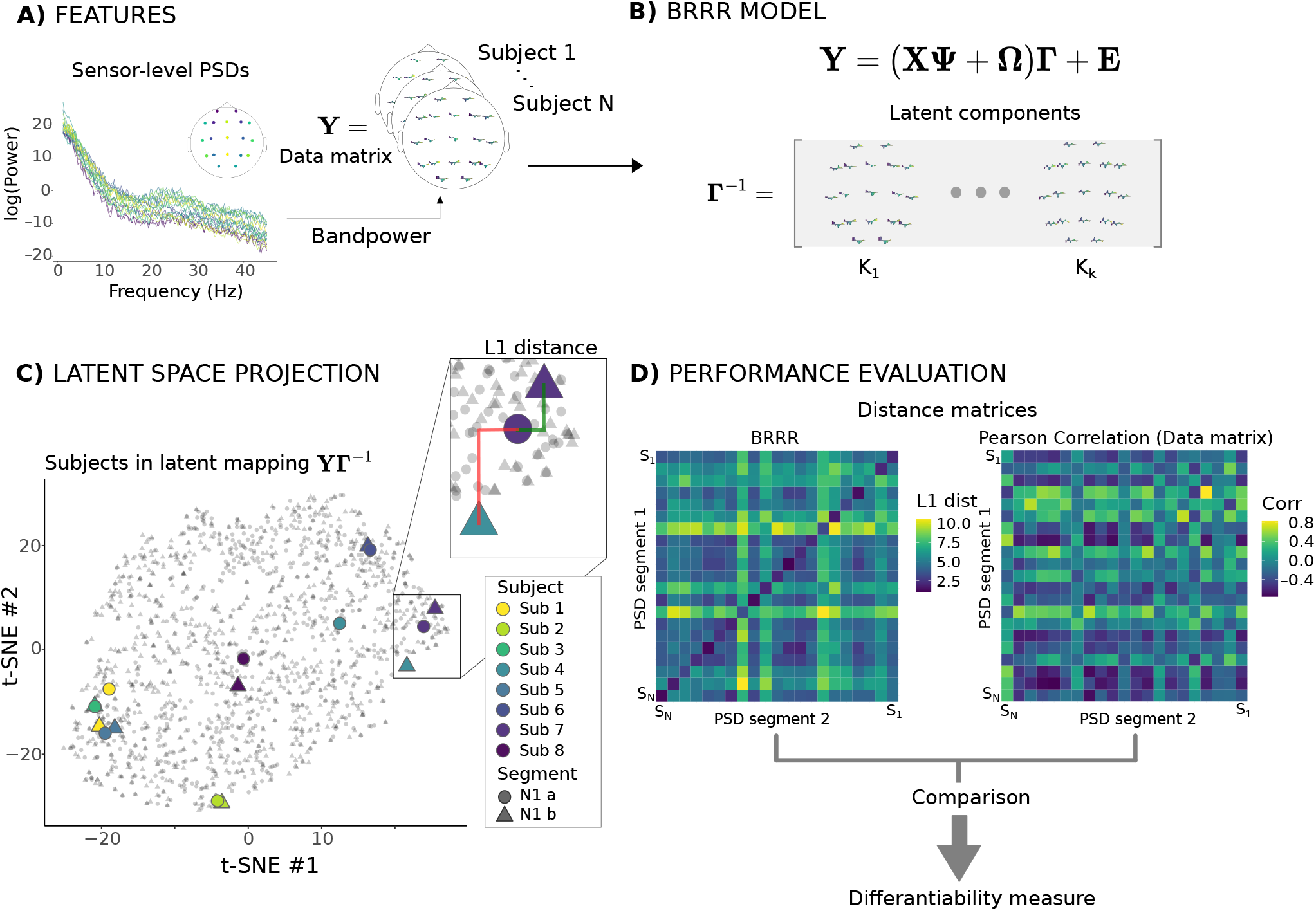
Overview of the analysis pipeline. We calculated, for each subject, the sensor-level PSD estimates (**A**) which were then vectorized and concatenated into a data matrix. The individual bandpowers in different sleep stages were used as observations in the BRRR model (**B**) which produces a low-dimensional representation that best differentiates between the subjects (here the loading matrix **Γ**). Each subject was then projected onto the low-dimensional space produced by BRRR (**C**), here visualized, for simplicity, in two dimensions using stochastic neighborhood embedding (tSNE). The distances between subjects’ own data projections (within-subject distance) were compared to the distances to other subjects (between-subject distances). These distances were collected into distance matrices (**D**) from which the self-distances were defined as the within-subject distance compared to the average between-subject distances. The distance matrix resulting from application of the BRRR model was further compared to a distance matrix based on Pearson correlation of the full-dimensional data matrix **Y**.

In the BRRR model, the regression equation for the measured EEG data (see Figure 3B and equation 1 in Modeling), encodes the assumption that noise and covariates affect the response via a shared subspace **Γ** and approximates the response using a low-rank decomposition. The model aims at finding the loadings **Γ** that effectively maximize the distance between subjects while minimizing the within-subject distance (Figure 3C–D). For feature extraction, a low-dimensional representation of the data is useful. While most of the information from the original data is retained, the low-dimensional representation makes the task for the machine learning algorithm easier without sacrificing the model performance. Here a low-dimensional representation of the data was obtained by enforcing a latent space dimension smaller than both the number of features (*s*) and observations (*p*), i.e., *k* = 30 ≪ *p, s*. This is achieved by imposing shrinkage priors to **Ω** and **Γ** (see [24]).

To evaluate how well the model differentiated between subjects, we computed the L1 distances between test subjects’ EEG bandpower segments which were projected into the low-dimensional latent component space produced by the BRRR model. A subject was considered correctly identified if the minimum distance over any set of distances occurred between the individual’s own data segments (see Figure 3C). The success rate of the BRRR model was estimated using 10-fold cross validation (CV) and the percentage of correctly differentiated test subjects [21]. To examine the stability of the EEG fingerprint over the course of the sleep recording, we tested how well the BRRR model generalizes to subsequent PSD segments of the same individual. Ideally, the model should correctly predict to which individual a new observation belongs. For this, we used an additional out-of-sample 10-fold cross validation procedure, where the distances were measured between test subjects and an additional PSD-segment from N1 or N2 sleep, projected into the latent space.

### Within sleep stage models result in good fingerprinting accuracy

We first conducted the analysis within sleep stages. For both subject groups (‘All’ and ‘*>*7-year-olds’), test subjects could be differentiated with over 87% success rate (88% for All, 87% for *>* 7-year-olds) when spectra from two N2 segments were used as observations in the training, and with over 73% success rate when two N1 data segments were used (see Figure 4A; For more information, see Table 1 in Supplementary Tables). The N2 model explained up to 85% of the variation in the response (proportion of total variance explained, PTVE), indicating that most of the variation in the EEG bandpower within sleep stage can be attributed to the individuals. Adding a third N2 segment to the training data slightly increased the success rate for group ‘All’ (90%), while staying the same for ‘*>*7-year-olds’, again demonstrating good stability of the low-dimensional fingerprints. The model fit remained high at 80%.

**Table 1.** The 10-fold CV results of different BRRR models trained with K=30 components, tabulated for both within and mixed sleep stage models. ‘Input’ refers to the data used here for training the models. Success rate (SR) and proportion of total variance explained (PTVE) were averaged over the folds. The best results are highlighted. Column ‘SR corr’ marks the success rate of correlation-based subject differentiation. The BRRR models that differ significantly (*p <* 0.01) from correlation-based method are marked with an asterisk.

| Group | Input | All |  |  |  | >7 year olds |  |  |  |
| --- | --- | --- | --- | --- | --- | --- | --- | --- | --- |
|  |  | SR | PTVE | SR Corr | . | SR | PTVE | SR Corr | . |
| Within sleep stage | **2 N1** | 0.73 | 0.77 | 0.60 | * | **0.89** | **0.78** | **0.78** | * |
|  | **2 N2** | **0.88** | **0.85** | **0.82** | * | 0.87 | 0.82 | 0.83 | ns. |
|  | 3 N2 | 0.90 | 0.80 | 0.82 | ns. | 0.87 | 0.75 | 0.86 | ns. |
| Mixed sleep stage | N1 + N2 | 0.24 | 0.60 | 0.15 | * | 0.44 | 0.57 | 0.29 | * |
|  | **2N1 + N2** | **0.72** | **0.64** | **0.60** | * | **0.90** | **0.61** | **0.78** | * |
|  | N1+ 2N2 | 0.29 | 0.60 | 0.19 | * | 0.54 | 0.56 | 0.37 | * |

**Figure 4.**
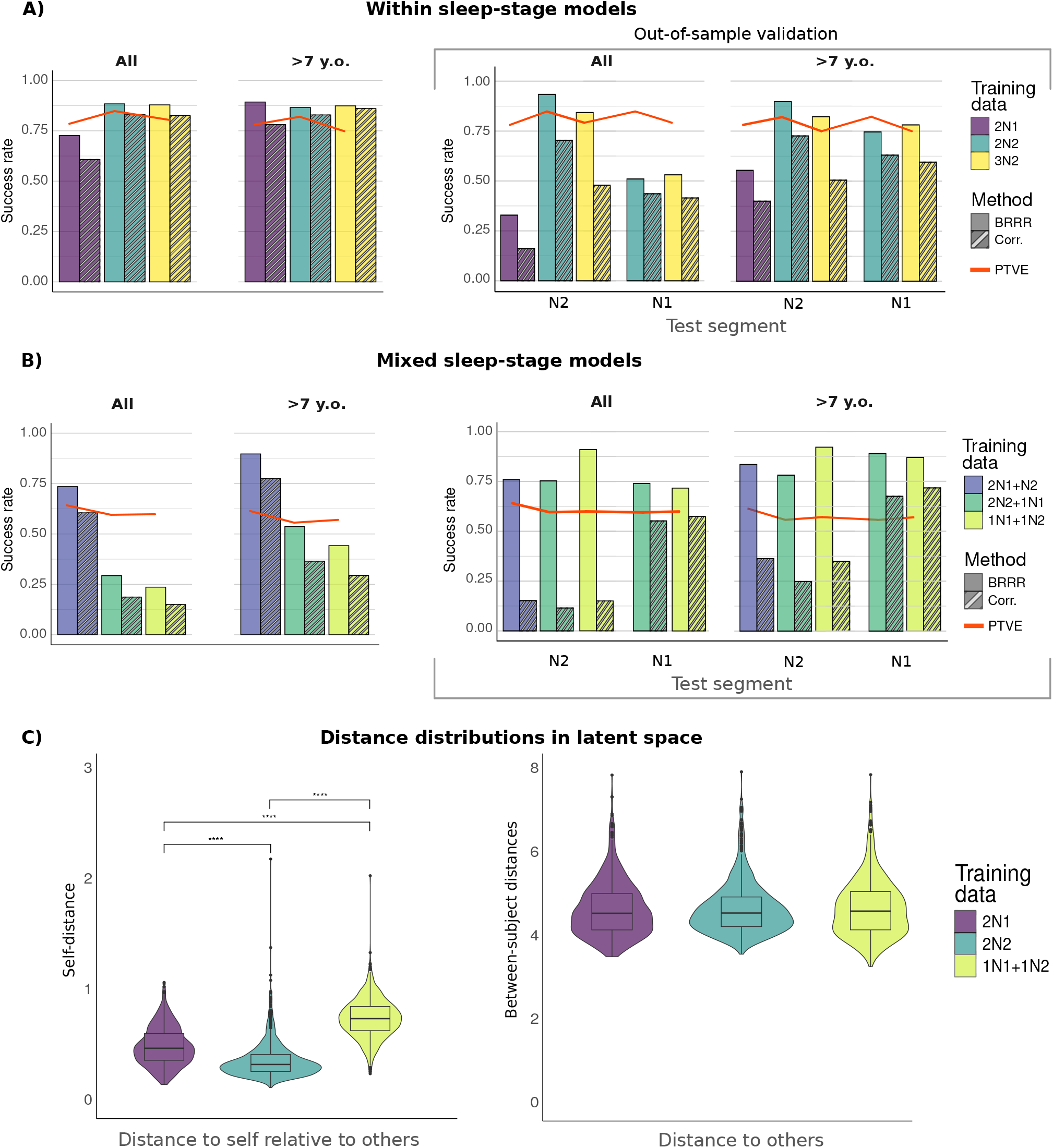
Differentiability scores across different models and subject groups. **A**: Averaged 10-fold cross-validation success rates for the within-sleep stage models. Left: Success rates for differentiating subjects in groups ‘All’ and ‘*>*7-year-olds’. The red lines indicate the proportion of the total variance (PTVE) explained by the model. Right: Corresponding Figure for both groups when predicting the identity of an additional test data segment. **B**: Averaged 10-fold CV success rates for mixed sleep stage models. Left: Differentiation of test subjects in both groups, Right: Prediction of the identity of an additional data segment. **C**: Distance distributions in the latent space (note the different maxima of y-axis). Left: the self-distance score distribution in two within-sleep stage models (2N1, 2N2) and one mixed sleep stage model (1N1+1N2). Bars with asterisks indicate significant differences between models. Right: Average distance distributions to others using the same models.

For the N1 sleep, the models trained with data from over 7-year-olds performed better than the models trained with all data: using two N1 data segments as training data yielded differentiability scores 89% and 73%, respectively (Figure 4A and Table 1 in Supplementary Tables).

We then examined how well the model trained within one sleep stage data generalized to a previously unseen observation either from the same or the other sleep stage (Out-of-sample validation: Figure 4A, right panel and Table 2 in Tables). Unsurprisingly, if the model is trained within N2 data, the latent space projection maps an additional N2 test segment close to the individual representation, resulting in success rates of up to 92% for ‘All’ and 90% for ‘*>*7-year-olds’. For ‘*>*7-year-olds’, the model trained within N2 sleep could also predict the identity of N1 test segment with good, 75-78% success rate, indicating good generalizability, but for the model trained on the full dataset, the success rates were lower (54–55%). For the inverse situation of predicting the identity of N2 test segment using the latent representation from N1 sleep model, the success rate decreased, resulting in lowest success rates for both groups (‘All’: 32% ‘*>*7-year-olds’: 55%).

**Table 2.** Summary of the different BRRR models using different inputs for across-stage prediction. Success rates (SR) and PTVE scores were averaged over 10 CV test samples. Here ‘Test’ refers to the sleep stage of the PSD segment used in the validation. The model with highest accuracy is emphasized. Significantly (*p <* 0.01) different models are marked with an asterisk.

| Group | Input | Test | All |  |  |  | >7 year olds |  |  |  |
| --- | --- | --- | --- | --- | --- | --- | --- | --- | --- | --- |
|  |  |  | SR | PTVE | SR Corr | . | SR | PTVE | SR Corr | . |
| Within sleep stage | 2 N1 | N2 | 0.32 | 0.79 | 0.16 | * | 0.55 | 0.78 | 0.40 | * |
|  | **2 N2** | **N2** | **0.92** | **0.85** | **0.69** | * | **0.90** | **0.82** | **0.73** | * |
|  | 3 N2 | N2 | 0.87 | 0.80 | 0.47 | * | 0.87 | 0.75 | 0.49 | * |
|  | 2 N2 | N1 | 0.54 | 0.85 | 0.43 | * | 0.75 | 0.82 | 0.63 | * |
|  | 3 N2 | N1 | 0.55 | 0.80 | 0.41 | * | 0.78 | 0.75 | 0.60 | * |
| Mixed sleep stage | N1 + N2 | N2 | 0.70 | 0.60 | 0.58 | * | 0.87 | 0.57 | 0.72 | * |
|  | 2N1 + N2 | N2 | 0.76 | 0.64 | 0.16 | * | 0.84 | 0.61 | 0.36 | * |
|  | **N1 + N2** | **N1** | **0.90** | **0.60** | **0.17** | * | **0.92** | **0.57** | **0.35** | * |
|  | N1+ 2N2 | N1 | 0.76 | 0.60 | 0.55 | * | 0.89 | 0.56 | 0.68 | * |

### Mixed sleep stage models produce more generalizable fingerprints

In the next step, we trained the models by using segments from both sleep stages: the results are summarized in Figure 4B and in Tables 1 and 2 (Supplementary Tables). When the training data contained two N1 data segments and one N2 segment, nearly 90% success rate was reached for ‘*>*7-year-olds’, and 72% success rate for the full dataset. However, the success rate for subject differentiation dropped below 30% for ‘All’ when the training data included only one N1 data segment combined with one or two N2 data segments, probably related to more heterogeneous input data. For ‘*>*7-year-olds’, the success rates were higher (44–54%) than for models trained on the full dataset, although clearly lower than within-sleep stage success rates. Although the additional variance associated with sleep stage properties could no longer be predominantly attributed to individual differences (as indicated by the decrease in the model fit), the BRRR model still accounted for most of the variation in the response (PTVE *>*56%).

The model success rates increased when using the out-of-sample validation procedure: for example, when using one segment from each of the sleep stages in training, the model success rate for N2 test data was 90% for group ‘All’ and 92% for ‘*>*7-year-olds’, the latter exceeding the results for within sleep-stage fingerprinting (Figure 4B, right panel). When testing the same model on N1 data segment, the success rates were at 70% (‘All’) to 87% (‘*>*7-year-olds’).

Finally, using three segments in training, two of them from the same sleep stage, produced generalizable fingerprints for both groups: depending on the test data segment, the success rates were about 76% (‘All’) and 78–89% (‘*>*7-year-olds’). Overall, the participants in the older age group seemed to have slightly better results within and across sleep stages, thus potentially having more reliable within-session sleep fingerprint.

### Stability of the low-dimensional representation is affected by age

As suggested by the within N1 sleep and mixed sleep stage results above, the stability of individual features across sleep stages appeared somewhat age dependent. Indeed, age emerged as an organizing factor in the latent space learned from N1 data segments, as illustrated in the two-dimensional t-distributed stochastic neighborhood embedding (t-SNE; [60]) map in Figure 5. The sex of the participants, however, did not similarly separate the subjects in the latent space.

**Figure 5.**
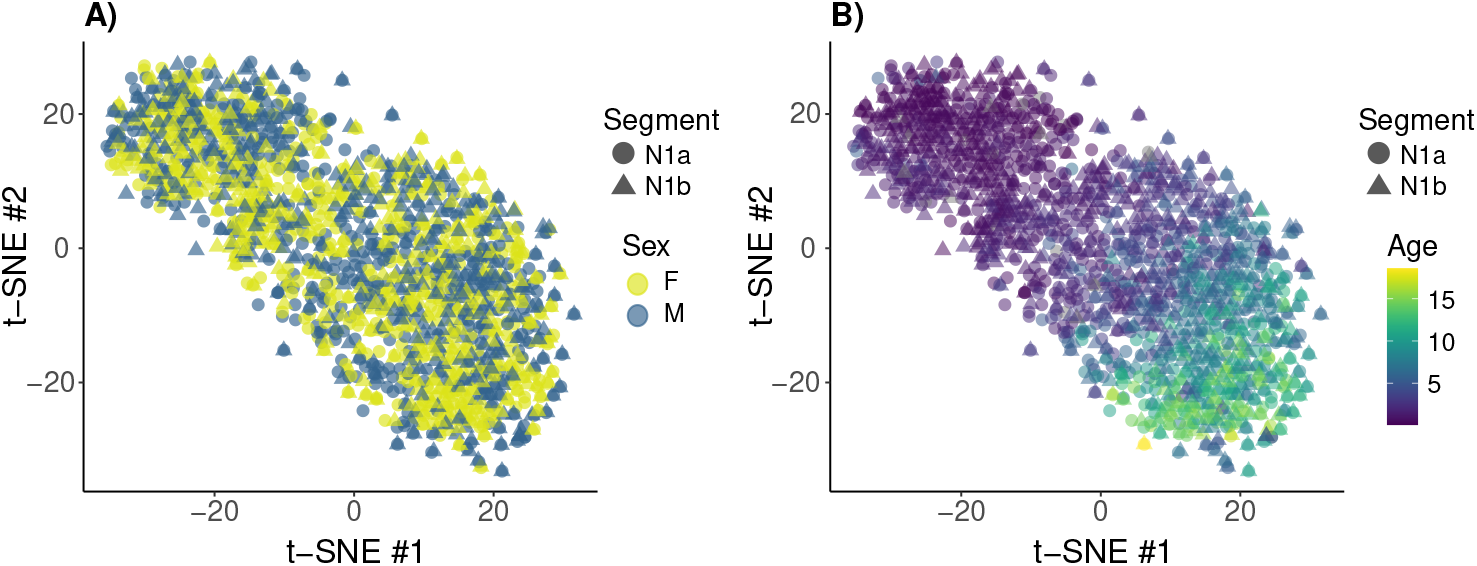
Two-dimensional projection of the subjects’ latent spaces (2N1 model trained with one N1 data segment and tested on another N1 data segment), coloured by sex (**A**) and age (**B**) and visualized with t-distributed stochastic neighbor embedding. The first N1 segment is marked with a circle, the second with a triangle. The age of the participants emerges as an organizing structure in the latent space while the sex does not.

As a measure of stability of the latent representation, we utilized self-distance, defined as the ratio of distance to self, divided by the average distance to others (see Eq. 3 in Statistical testing). Here, statistically significant differences in the self-distance scores between the sexes emerged for N1 sleep data (*p <* 0.001) but not for N2 (*p* = 0.94). In mixed sleep stage latent space representation, containing data from both N1 and N2 sleep, there was a small but significant difference between sexes (*μ*_self-diff._*F* = 0.74, *μ*_self-diff._*M* = 0.77, *p* = 0.042).

Self-distance scores were affected by the combination of training and test data segments used in the model (Welch ANOVA across the means of self-distance distributions, *p <* 0.001, Figure 4C). In line with the results above, the self-distances were lowest for the representation learned from two subsequent segments of N2 sleep data, highest in the mixed sleep models, and in between for N1 sleep. However, the distributions of average distances to others did not seem to change with the training data (4C, right), suggesting inter-subject variance of similar magnitude regardless of the sleep stage.

Since we could not fully exclude the possibility that the self-distance measures in the latent projection are influenced by gender to some extent, we performed a regression analysis to study the effect of age separately for boys and girls. Figure 6 illustrates the regression fits for the self-distance scores for N1 sleep and mixed sleep representations as a function of age, separately for girls and boys. For girls, age explains 7.3–8.2% of the variation in the self-distance scores (N1 sleep and mixed sleep models, *p <* 0.0001). For boys, age explains 2.1–6.3% of the variation (N1 sleep and mixed sleep models, *p <* 0.005). For both sexes, the self-distance measures decreased as a function of age, indicating increasing stability.

**Figure 6.**
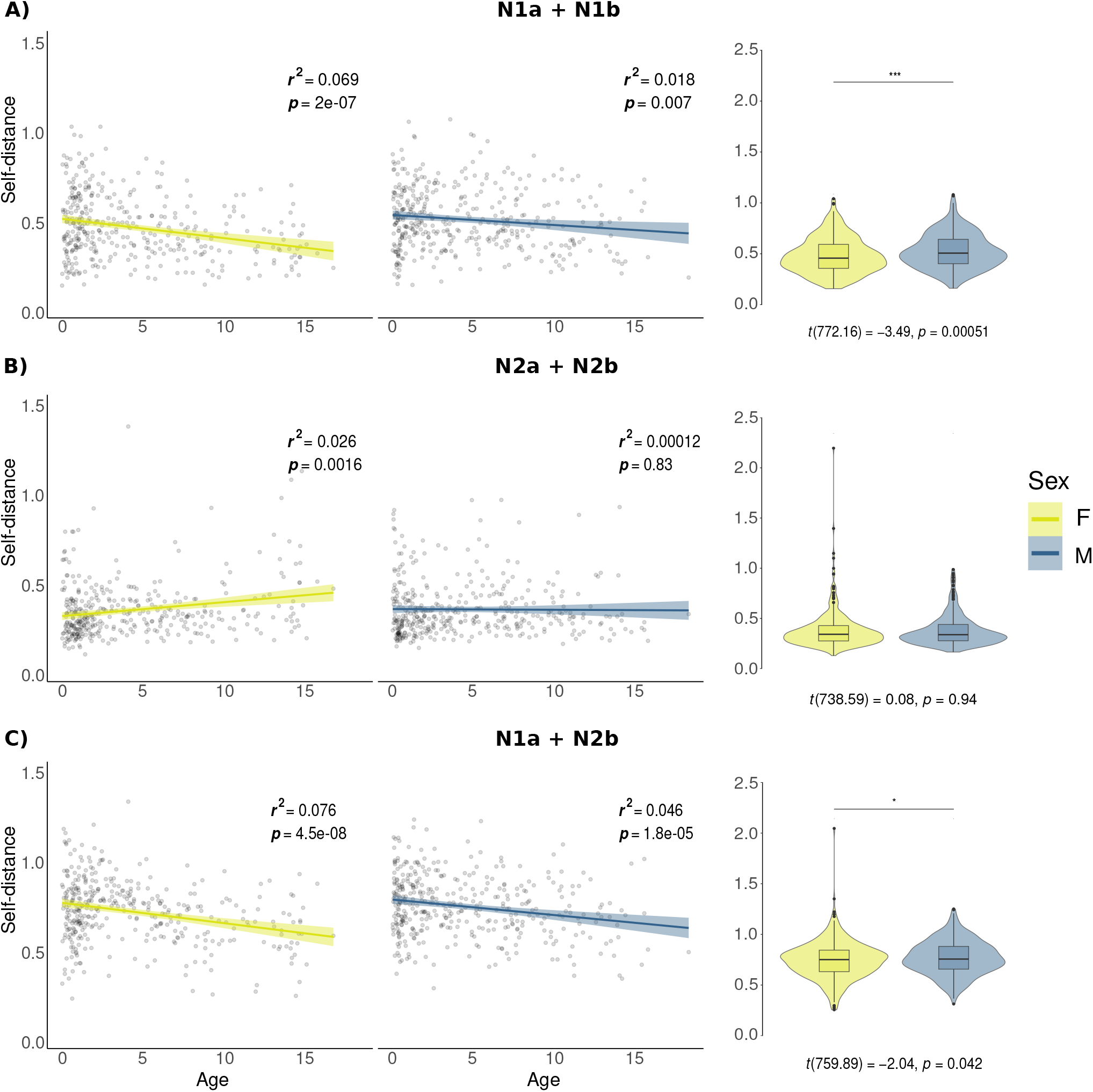
Linear relationship between age and self-distance (Left: shaded area represents 95% confidence intervals) and self-distance distributions (Right: mean comparisons are done with t-test) in the latent space over the whole population, separately for boys (M) and girls (F). Upper Figure **A** has self-distance scores within N1 sleep, middle Figure **B** contains the self-distances within N2 sleep and lower Figure **C** has the same scores in mixed (N1a and N2b) sleep latent space representations.

Age appeared as a relevant factor affecting the self-stability measures. In the N2 sleep model, a significant inverse relationship with age was observed for girls (self-distances increased as a function of age, *r*^2^ = 0.026, *p* = 0.0016), but not for boys, for whom age explained less than 0.1% of the variation in self-distance scores (p = 0.54; Figure 6 B). Notably, the girls’ N2 self-distance distribution contained several outliers. Furthermore, the learned representations were increasingly alike across sleep stages when participants aged: in the older subjects, the latent representations derived from the models trained within N1 or N2 sleep were more similar to each other (Mantel’s *r* = 0.523, *p <* 0.001) than models trained with all the subjects (Group ‘All’, *r* = 0.464, *p <* 0.001) or under seven-year old subjects (*r* = 0.412, *p <* 0.001).

### BRRR fingerprints generalize better compared to correlation-based fingerprinting

The BRRR model was bench-marked against a correlation-based similarity measure, which is commonly used in functional brain-fingerprinting [2, 11, 49]. Unlike in BRRR, the correlations between observations were computed on the full (test) data matrix **Y**, without any dimensionality reduction. The success rate was determined by averaging over the same 10 sets of test subjects and their corresponding data segments as for BRRR in the 10-fold cross-validation; when the number of observations per subject exceeded two, the maximum correlation was chosen across the test segment and the remaining observations. The results are summarized in Figure 4. Within N2 sleep, the BRRR and the correlation-based null model had comparable performances: the success rates for the model trained with 3 N2 segments did not differ significantly between BRRR and correlation-based fingerprinting for either of the groups (*p*¿0.05). The correlation metric provided the highest success rates when using data from N2 sleep only (average success rates between 83%–86% for both groups ‘All’ and ‘*>*7-year-olds’; see Figure 4A). Interestingly, the correlation-based model prediction seemed to suffer more from the increase of observations within sleep stages (4 N2 segments, of which one was used for testing), with success rate dropping from 70% to 48% for group ‘All’ (Table 2 in Tables), suggesting that the latent space projection might provide more stable individual representation within sleep stages. Similarly to BRRR, the correlational fingerprint showed generally improved success rates for the older group.

We also compared the performance of correlation and BRRR in the mixed sleep stage settings, where BRRR outperformed the correlation-based subject differentiation in all data conditions (*p <* 0.05, Figure 4B). For group ‘*>*7-year-olds’, BRRR was able to predict which new observation belonged to whom with up to 92% success (see Table 2) when trained with data segments from both sleep stages, clearly outperforming the correlation-based model (prediction success rate 35%). Group ‘All’ had corresponding results. Thus, a similarity metric based on just correlation does not appear to generalize across sleep stages and suffers from increased dimensionality and heterogeneity of the training data.

## Discussion

Our individually unique brain structures allow for identification based on anatomical brain features [58], much like the idiosyncratic patterns in fingertips. In recent years, also functional fingerprinting based on individual variation, applied both in M/EEG and fMRI, has gained momentum: especially in adult populations, features like functional narrow-band and broad-band connectome and power spectra have demonstrated fingerprint-like qualities when comparing to reference data from the same individual [6, 11, 27, 36]. In this study, we examined the possibility of EEG-based fingerprinting in a large pediatric population (N=782) and assessed the stability of broad-band power features within and across N1 and N2 non-REM sleep stages.

### Individual fingerprints generalize across sleep stages

Exploiting the correlation structures in the EEG broad-band power using BRRR yielded a latent-variable representation of the data that could differentiate the individuals from each other and predict identity with up to 93% average success rate. Furthermore, the individuals explained most of the variance in sleep EEG bandpower in the different modeling settings.

The models trained within one sleep stage had the highest success rates. Unsurprisingly, the explanatory power of the BRRR approach, measured by PTVE values, was highest for these models where non-individual related variance was not increased by adding data segments from different physiological states. Still, the model could fit into individual data very well across sleep stages, demonstrating the robustness of BRRR in conditions with different noise.

Despite the different spectral properties of N1 and N2 sleep stages, the BRRR algorithm could find a latent representation from their combination that allowed for excellent identity prediction up to 92% from subsequent data segments. However, the prediction success rates were generally lower for the mixed sleep-stage models. The relative self-distances in the latent space mirrored the model success rates, being lowest for model trained within N2 sleep and highest for mixed sleep stage model. We did not achieve as high fingerprinting accuracy scores as reported in some studies using PSD features extracted from MEG data [11, 27, 36]. This might be due to low-density EEG used here with relatively granular bandpower features, which may fail to capture all dynamic fluctuations in the brain oscillations. Also, the infant-skewed age distribution might lower the fingerprinting accuracies due to physical and functional maturation effects discussed in more detail below.

We observed systematic differences in the power spectral densities both between and within the sleep stages: for example, the power in the sleep spindle range (12–15 Hz) increased within N2 sleep across subsequent segments. The differences within sleep stage highlight that in longer sleep recordings, fingerprinting approaches within one sleep stage do not necessarily work. Given the non-linear nature of change in total power levels and considerable individual and sensor dependent variation in the oscillatory power, related e.g., to cap placement, skull thickness and cranial ossification, we opted to use relative bandpower measures over absolute bandpower. This does not, however, erase all the systemic differences in oscillatory power that were observed within and between the segments, like power variations at the typical sleep spindle frequency range.

### Age affects the differentiability of subjects

We had hypothesized that the stability of the fingerprint-like latent variable representations would increase as children mature. Indeed, in demanding settings, such as when differentiating individuals across sleep stages or predicting the identity of a test data segment using a representation learned from another sleep stage, concentrating the analysis to older participants led to generally better performance in terms of success rates, both for BRRR and the correlation-based model. When comparing the latent representations of the within-sleep stage models, the representations learned from over seven-year-olds were more alike than those learned from all the subjects or under seven-year-olds. The representations were thus more generalizable in the older subjects, allowing across sleep stage prediction. Furthermore, we observed age-related linear trends in the self-differentiability scores. In our analysis, representations learned from N1 sleep and the mixture of N1 and N2 sleep representations showed the most prominent age effects. The relative bandpower estimates used here contain also aperiodic signal parts [18, 57], which have been demonstrated to be both individual [15, 44] and subject to changes during maturation [57]. It is therefore possible that the age-effects observed in the low-dimensional space reflect changes particularly in the aperiodic power. These may include changes related to skull-thickening and ossification during the first year of life. Furthermore, flattening of the aperiodic activity and slowing of alpha oscillations continue across the human lifespan [43]. Possible age-related differences in sleep behavior may also confound these results, especially in mixed sleep stage model where the PSD segments are further apart from each other in time.

We observed small but significant sex effects in the self-distance scores for N1 and mixed sleep stage models, but not for N2 model. In a longitudinal adolescent population study, Candaleria-Cook et al. reported significant age-related changes in relative EEG power specifically in males [7].

Instead, in our study the self-distance scores for females slightly increased in N2 sleep as a function of age, indicating decreasing stability, but no significant relationship was observed for males. In this sleep stage, the self-distances were generally very small across all the subjects, thus making the regression fit more sensitive to outliers. Furthermore, the skewed age distribution increases the uncertainty of the regression fits in the older age brackets. Possible differences between sexes may also be attributable to external factors, such as non-equal distribution of movement artifacts.

### Benefit of latent variable approach in modeling individual differences

Earlier results on the usefulness of data-reduction techniques for fingerprinting have been mixed, some reporting no improvement in fingerprinting success rates [11] while others claim considerable benefit [2, 34]. As opposed to performing subject differentiation based on full feature matrices, we opted to use a latent variable approach, leaning into the intuition that not all features are informative nor independent due to spatial correlations in the fingerprinting task [36]. Furthermore, using the individual identifiers as variables helps to ensure that the low-dimensional space maintains features informative of individuality, as opposed to unsupervised dimensionality reduction methods.

In almost all cases, barring the models trained exclusively within N2 sleep, subject differentiation in 30-dimensional latent space provided by BRRR significantly outperformed the correlation-based differentiation on full 247-dimensional feature matrix. The differences in these two methods were most notable when trying to predict the identity of an additional data segment across sleep stages: for example, when including data from both sleep stages in training and using N1 sleep as the test segment, the difference between BRRR and correlation-based fingerprinting accuracy scores was over 50%. This is likely due to the low-dimensional representation of the data which succeeds in reducing the noise (including measurement noise and non-modeled effects) while refining the individual features. Thus, latent mapping can maintain low within-subject distances even when presented with new data. A previous study using BRRR to source-localized MEG found slightly better results with correlation-based fingerprinting than reduced-rank regression [27]: we suggest that in the case of large sample-size sensor-level clinical EEG, the latent noise model is especially beneficial.

Fingerprinting accuracy benefited from informed regularization: the BRRR approach did not suffer as much from the increased number of subjects as the correlation-based model. However, the BRRR model was sensitive to overfitting: including too many components reduced the model’s fingerprinting performance drastically while the amount of explained variance by the model stayed stable.

### Fingerprints over developmental periods

To establish a long-term EEG fingerprint, the BRRR model should generalize to new observations from the same individuals, preferably over separate measurement sessions, experimental tasks, or states. Our results point to stable functional fingerprints in developing children across sleep stages measured within one session, but long-term fingerprints over separate sessions measured far apart have not yet been established.

The short-term test-retest reliability (*>* 1 week) in M/EEG oscillatory activity seems to be good in children [37], but maturation changes the oscillatory activity patterns significantly in the long term in school-aged children [7]. As infants undergo even greater development during their first year of life, including the emergence of rhythmic oscillatory activity [41, 48], the functional dynamics in this stage are less likely to preserve into adulthood. Interestingly, the maturation-related changes in the power spectra seem to be very consistent across individuals during the first two years of life, after which individual differences start to emerge [48]. Rapid physical changes during development, such as skull thickening and head size growth, also affect the conductivity and thus, the EEG features over time. While the functional fingerprints may not generalize over the developmental period within individuals, such a behavior might even be a characteristic feature indicating typical development: if maturation occurs normally, the power-spectral fingerprints identified in infancy should fade after certain (yet to be specified) time. Indeed, the short-term reliability in EEG-PSD estimates tends to be higher in typically developing children compared to children with autism spectrum disorder [37].

With a large children cohort such as the one used in this study, normative modeling [41], akin to growth charts, could be utilized for comparing pediatric patients with developmental conditions to typically developing children. For example, abnormal sleep EEG patterns, such as those present in epilepsy, might manifest in the inter- and intra-subject distances in the latent space projection across and within the sleep stages. Conversely, successful treatment of clinical conditions might manifest as improvement in the within-subject reliability estimates. Building normative models using latent variable formulation might be especially useful in settings where the data is more heterogeneous or of variable quality. Furthermore, they could allow modeling associations between functional neuroimaging data and behavioral scores beyond correlational inferences [9] while accounting for between-subject variance. Getting representative subject samples with low-density sleep EEG is cheaper and easier compared to other functional neuroimaging modalities, EEG being often the only feasible brain imaging modality to collect data from children. Thus, normative pediatric sleep-EEG models could be of great clinical importance in the future. For this purpose, distance-based fingerprinting measures appear to be especially useful.

### Limitations of the study and outlook

The EEG data used in this study was preprocessed only lightly to ensure equal amount of data for all the participants. Therefore, we did not, e.g., remove noisy segments or use automated artifact rejection algorithms. Independent component analysis was applied to remove the heartbeat artefact, and bandpass filtering took out most of the slow drifts, but some muscular and ocular artefacts likely remained in the data. Any discrepancy in artefact density, such as uneven amount of movement between sexes in N1 sleep, might account for some of the observed sex differences. Also, the sleep segments were labeled as “N1” or “N2” sleep based on the emergence of first K-complex alone, but it is likely that the segments contained data from other sleep stages or wakefulness. Furthermore, it is possible that some of the latent components of the BRRR model reflect non-brain activity. Our work was restricted to low-density sensor-level analysis: using source-level features instead might allow to investigate which brain areas contribute to individual differentiability and how they might change across age, in addition to providing better anatomical accuracy overall. To perform reliable source-level modeling, a higher-density electrode cap of preferably at least 64 electrodes [29] and structural MRIs would be needed. The same holds true for conducting connectivity analyses, which are commonly used in fingerprinting to produce highly individual networks. In addition, differences in EEG cap placement may have considerable effects for frequency topographies used in fingerprinting. This should be tested in longitudinal data sets with recording sessions conducted in different days.

Even though most frequency-based features used in subject differentiation are derived from oscillatory power in pre-defined frequency bands, such as theta, alpha, beta or gamma bands, considerable individual dynamics of the aperiodic component of PSDs have been reported as well [15, 17]. EEG fingerprinting based on the aperiodic component of the power spectra, in addition to broadband power, was reported to have high (95%) identification accuracy in the adult population [15]. Developmental and maturation-related trends seem to be reflected in the aperiodic power as well: the oscillatory activity of infants is mostly aperiodic [50], which in the context of our study can mitigate successful across sleep-stage fingerprinting of that cohort.

An important caveat in the interpretation of the PSD-related maturational phenomena relates to the difficulty in determining whether, e.g., the decrease of alpha power during maturation relates to change in alpha amplitude or the incidence frequency of alpha oscillations, especially when using relatively long data segments for estimating PSDs [18], or to physical changes not related to functional maturation. Addressing the oscillatory activity in the temporal domain and in a data-driven manner with, e.g., hidden Markov models [62] could further clarify the individual and age-related dynamics in sleep stages. Another aspect to consider is the possible optimal data time windows for fingerprint estimation, which may vary based on the features used [59]. For resting state, reliable spectral estimates can be achieved from just 30–120 seconds of data [64], but, as demonstrated here, these are subject to changes during the progression of sleep.

## Conclusions

To conclude, we were able to differentiate participants from each other within one recording session with excellent accuracy based on the bandpower estimates of low-density clinical sleep-EEG. The differentiation result held even across sleep stages in a large cohort of children. We found an age effect in the self-differentiability measures of N1 and a mix of N1 and N2 representations, where the differentiability improved as a function of age. Using a probabilistic latent regression model was beneficial for the task, as our model outperformed the commonly used correlation-based differentiation. Within-session sleep fingerprints extracted by latent variable models may help to assess typical development in a normative manner. Further longitudinal research is needed for establishing the presence and evolution of functional brain fingerprints over developmental periods.

## Materials and Methods

### Subjects

The dataset contained sleep EEG recordings from 821 healthy Finnish children. The recordings had been originally collected due to clinical concern, but no neurological or developmental diagnoses had been given to the patients within four years after the EEG. The data were measured at Helsinki University Hospital, and the Institutional Research Review Board at Helsinki and Uusimaa Hospital district approved the study including waiver of consent due to the retrospective collection of data acquired as part of standard of care. The age of the children varied between 6 weeks and 19 years (mean ± SD age 4.6 ± 4.3 years) with uneven distribution: over half of the participants were infants (see Figure 7). Of the original data sample, 405 (49.3%) were females, 414 (50.4%) were males and 2 were not assigned sex. The sex ratios were mostly even across the age bins. After removing corrupted or too short data files (N=39), 782 recordings remained for analysis.

**Figure 7.**
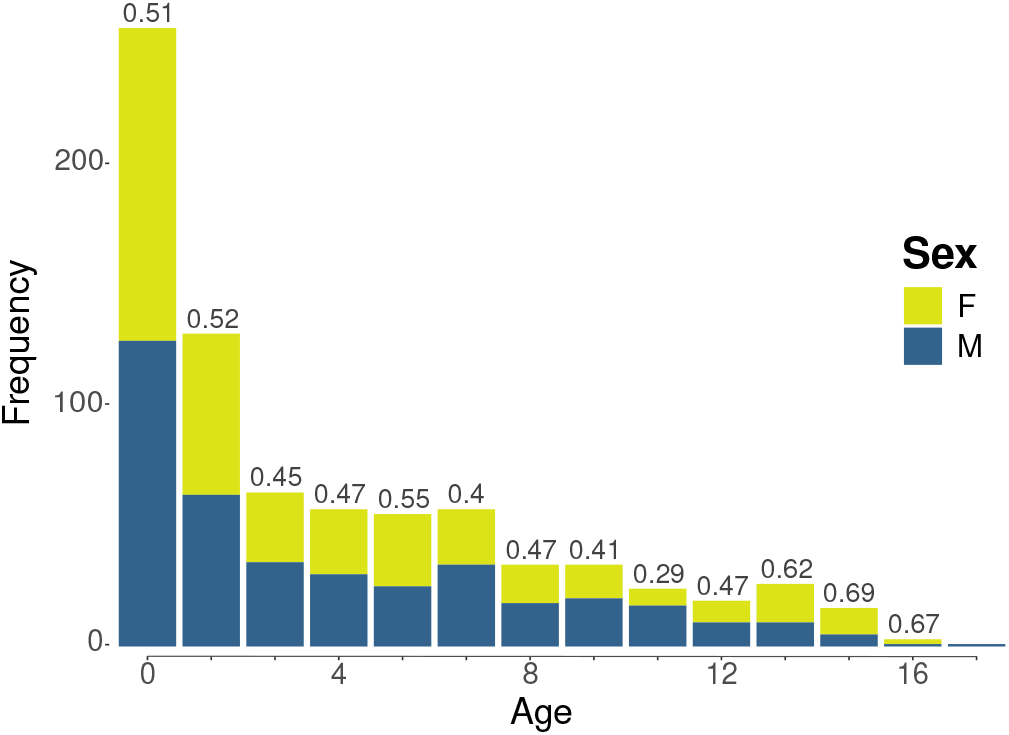
The age distribution of the subjects. The proportion of females in each age bin is marked on top of each bar.

### EEG recordings

The EEG sensors were placed using the standard 10-20 montage, and the data was referenced either to Cz or Pz channel. Eye movements were recorded with piezoelectric sensors. In addition, the recordings contained electrocardiogram (ECG) and breathing sensors. The subjects were measured using two different-sized caps; generally, children under the age of 5 were measured with a 19-channel cap which lacked one electrode compared to the one used in older children.

The EEG data segments used in the analysis spanned over 900 seconds. The first sleep spindle or K-complex [19], considered to mark the initiation of N2 sleep, was labeled manually by an expert technologist. The preceding 300 seconds were labeled as N1 sleep in the analysis, although from the raw data it was apparent that some of the subjects were periodically awake. The following 600 seconds were categorized as “N2 sleep”, although it is possible that the child was in lighter sleep or awake during this stage.

### EEG preprocessing

Preprocessing of the EEG data was carried out with MNEpython v1.5.1. To have an equal amount of data from each subject, the analysis was limited to EEG channels shared by every recording, totaling 19. All the data was re-referenced to average reference for homogenization.

The EEG time series were bandpass filtered at 1–45 Hz. After filtering, we used independent component analysis (fast-ICA algorithm, [30], implemented in MNE-Python [25]) to project out the prominent ocular and cardiac artifacts. To mimic typical clinical EEG analysis, no additional automatic artifact removal was conducted. All the data was manually checked for larger artifacts. Especially during the N2 stage, the data from the sleeping children rarely contained any blinks, and some of the muscle and movement artifacts were attenuated by the bandpass filtering applied.

Spectral estimates were calculated for each subject after data cleaning. The power spectral densities were calculated separately for both sleep stages (N1 and N2). The EEG time series was cut to 60 second epochs with inter-epoch intervals of 30 seconds. One-minute data segments are generally considered enough for estimating the spectral properties of both rhythmic and arrhythmic phenomena [36, 64]. We estimated the power spectral densities (PSDs) for a total of six epochs, 2 for N1 sleep stage and 4 for N2 sleep. More N2 epochs were chosen since the recordings contained twice as much N2 sleep compared to N1 sleep. The PSDs were calculated using Welch’s method with Hamming window (*n*_fft_ = 1024, sampling frequency = 250 Hz, frequency resolution ≈ 0.244) and averaging over the epochs. Examples of the PSD segments within subjects (in log-scale) are depicted in Figure 1C.

For the regression analysis, 13 separate frequency bands were defined, for which the relative power for each subject was calculated. The width of the frequency bands (1–3 Hz, 3–5.2 Hz, 5.2–7.6 Hz, 7.6–10.2 Hz, 10.2–13 Hz, 13–16 Hz, 16–19.2 Hz, 19.2–22.6 Hz, 22.6–26.2 Hz, 26.2–30 Hz, 30–34 Hz, 34–38.2 Hz, 38.2–42.6 Hz) was set to increase by 0.2 Hz in each consecutive band. To inspect global, age-related changes in the power spectral densities, the subjects were divided into 10 age groups of equal size (refer to the color bar in Figure 1A).

### Modeling

Data analysis was conducted with R v4.4 [47] and MNE python [25]. For an outline of the BRRR analysis process, see Figure 3. We aimed at finding latent spatio-spectral components that would maximally explain the variance between subjects while minimizing the distances between subjects.

The generative BRRR model was built as follows:

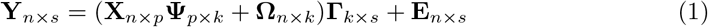

The response matrix **Y**_*n×s*_ contained either two or three PSDs over the 19 channels per subject (here *s* = 19 channels × 13 frequency bands = 247). Hence there were 2–3 observations per subject when training the model. The covariate matrix **X**_*n×p*_ was filled with 0–1 -entries, where the observations belonging to the same subject (indicated by the columns) were marked with 1, other entries being zero.

Essentially, **X** encodes which PSDs belong to which subject. **Ω**_*n×k*_ carries the unknown latent factors, or components *k*, which are taken to model the latent noise. The low-rank regression coefficient matrix **Ψ**_*p×k*_, together with projection matrix **Γ**_*k×s*_, (which projects latent space to EEG data, or to the observational space) constitute the standard regression coefficient matrix **Θ** = **ΨΓ**. Finally, **E**_*N×S*_, *e*_*i*_ ∼ 𝒩 (0, Σ) corresponds to unstructured and independent noise in the observational space. The noise matrix specified in Eq. 1 thus contains not only the residual noise, but also the differences within the participant’s own data.

The model fit is assessed with the proportion of total variance explained by rank *K* BRRR solution, excluding the latent noise. PTVE for the BRRR is defined as in [42]:

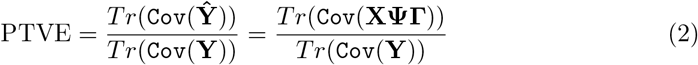

Where *Tr* is short for trace operation.

Conceptually, the BRRR model can be thought of as a combination of factor analysis and regression modeling. In the Bayesian context, the regression coefficients are learned using probabilistic inference. Unlike more customary dimensionality reduction procedures like PCA, BRRR encourages similar correlation structure to both regression coefficients and target variables via shared projection **Γ**, thus taking into account both the response and covariates in the dimensionality reduction. The product **XΨ** constricts the solutions so that the latent factors, or components, are shared when the observations belong to the same subject. For more details of the BRRR model specifications, see [24] and [42].

### Model training and convergence

Prior to applying the model, the EEG feature matrix **Y** was z-scored to have zero mean and unit variance across the features. The latent noise variance prior 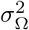 was adjusted to 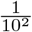. Thus, the latent noise levels are assumed to be relatively low while covariates are allowed to explain most of the variation. The noise priors of the model were tuned to suit the noise levels of EEG data; compared to priors set for MEG data in previous studies [27, 36], the expected amount of latent noise was set to be four orders of magnitude larger.

All BRRR models were initialized with Fisher’s linear discriminant (LDA) and their parameters were trained using the Gibbs sampler with 1000 iterations. The first 500 samples of the Markov chain Monte Carlo (MCMC) chains were removed as burn-in period, so that the parameter estimates were calculated from the last 500 samples. The convergence of the MCMC chains was assessed by computing the potential scale-reduction factors 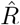 and bulk- & tail effective sample sizes (ESS) estimated for 200 randomly sampled indices in the regression coefficient matrix **Θ** = **ΨΓ**, analogously to [24]. The results, indicating good convergence and appropriate sampling of the posterior distribution, are summarized in Figure 8: the 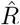 values are close to 1, and both tail- and bulk-essential sample sizes (ESS_*b*_, ESS_*t*_) are over 100, which are proposed rules-of-thumb for accepting the sample [23, 61].

**Figure 8.**
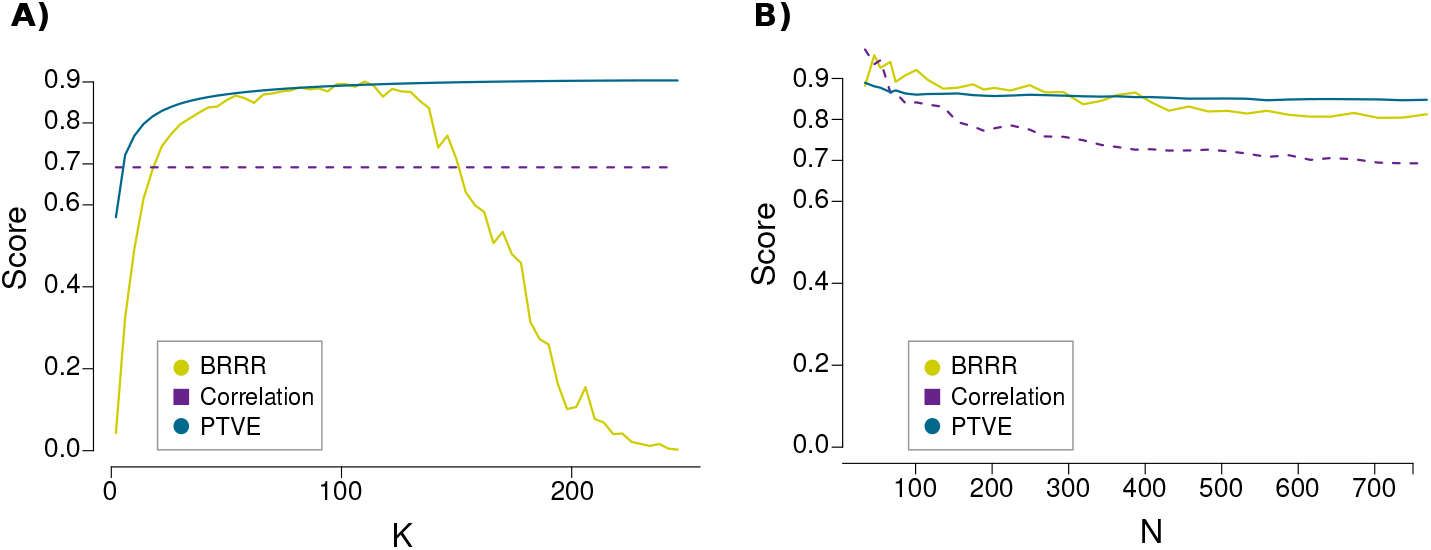
The model performance metrics (prediction success rate and proportion of total variance explained for BRRR) as a function of latent space dimension K (**A**) and number of subjects N (**B**), while keeping the latent space dimension fixed at *K* = 30. The models were trained with N2 sleep data: here the PTVE and success rate are estimated for the full data set, and both for the BRRR and the correlation-based null model. As the correlation-based method uses the full feature space, it does not have K-dependency (**A**).

In addition, model convergence was also reviewed visually by inspecting posterior PTVE traces (Figure 10 in the Supplementary Figures). All the checks suggest that 1000 iterations were enough for the Gibbs sampler to reach convergence. For all the convergence checks described here, we used an all-subject model with two segments of N2 data per subject.

### Performance estimation

#### BRRR

The input data from subjects were projected into the n×k dimensional latent space **YΓ**^−1^ inferred by BRRR. To assess how well the latent space structure managed to differentiate between participants, the within-subject distances were compared to between-subject distances and stored into a distance matrix. The minimum L1 distances were then used to determine the model’s success rates: the predicted identity of the test subject was the column index where the minimum distance was found. The choice of L1 distance as opposed to Euclidean distance was motivated by its better suitability for high-dimensional data [1].

#### Null model

For bench-marking our BRRR-fingerprinting results, we used a correlation-based “null model”. Similarly to [11], the distance matrix was calculated using Pearson correlations between observations. The predicted identity was assigned to the column index where the maximum correlation value was found. For this approach, we used all the features in the data without any prior dimensionality reduction. Pearson correlation was opted for as a similarity measure due to its common use in M/EEG fingerprinting.

#### Effects of feature and observation space sizes

We examined the BRRR model identity prediction performance as a function of latent space dimension K and number of participants. Here, the predictions were carried out within N2 sleep stage for the whole data set (782 subjects). The results are summarized in Figure 8A. The performance of the BRRR model had an inverse U-like relationship with the latent space dimension: the prediction success rate increased rapidly up to 30 components, reached its maximum of 90% around *K* = 100 and started to decrease rapidly after that, towards near zero at *K* = 250. The PTVE increased rapidly when K increased from 1 to around 50 but improved only marginally after that. The decreasing prediction performance allured to overfitting, as the latent space started to model noise that is not related to the individuality in the data. Between *K* = 20 − 150, BRRR had better prediction performance than the correlation -based method. Based on these investigations, we set the latent space dimension to *K* = 30 using the elbow method. The correlation-based null model performance was affected more by the number of randomly sampled participants than the BRRR model (Figure 8B): the correlation-based method seemed, however, to have an advantage over smaller (*N <* 75) sample sizes. The goodness-of-fit metric for the BRRR model, PTVE, was barely affected by the increasing number of participants after exceeding 100 subjects.

#### Cross-validation

For the subsequent analysis, model prediction accuracy (success rates) and training-PTVEs were estimated by averaging over 10-fold cross-validation results on test data (for success rate) and training data (training-PTVE estimates for model fit). We used two slightly different validation approaches. First, 10-fold cross validation was carried out on the test participant data using corresponding PSD segments that were used for the training subjects. The distances were computed by projecting the test subjects onto the latent space learned from the input data. To estimate the generalization ability of the BRRR model to new data segments, we used a slightly modified 10-fold cross validation scheme (out-of-sample validation). As previously, the test subjects were projected into the latent space, but now with an additional data segment from the same session that was not included in the training set. Next, we computed distance matrices by measuring the distances between the test subjects and these new data segments.

### Statistical testing

Cluster-level permutation tests [40] were used to determine if the PSD estimates of within N1 or N2 sleep stages differed from each other. The cluster-forming threshold was set to correspond to p-value of 0.01. The clusters were thresholded using an F-distribution with degrees of freedom *d*_1_ = 1, *d*_2_ = *n*_subj_ (N1 sleep) or *d*_1_ = 3, *d*_2_ = *n*_subj_ (N2 sleep). The analysis was carried out first for all the subjects (*n*_subj_ = 782), and then for the ten age groups separately (*n*_subj_ = 78). We also applied cluster-level permutation test for each age group to assess if the subjects’ averages over N1 and N2 segments differed from each other in the sleep spindle band (12–15 Hz). We used the same significance limit as before, and thresholded the clusters using an F-distribution with degrees of freedom *d*_1_ = 1, *d*_2_ = 78. To test the overall effect of age on the total power (see Figure 1), here determined as the area under the PSD curve (AUC), we fit a one-way ANOVA. For post-hoc analysis of the differences between mean AUC values of the age groups, a two-sided t-test was employed. To correct for multiple comparisons, we used Bonferroni correction for the p-values, which were deemed significant if they were under *p <* 0.0011 (significance value *α* = 0.05 divided by the total of 45 comparisons).

The difference in success rates, i.e., the correct data segment matching rate, of BRRR and correlation-based fingerprinting in the test data was statistically tested using a 10-fold cross-validated t-test [16]. We visualized the latent space projection of the subjects by using t-distributed stochastic neighborhood embeddings [60] (Figure 5), which is a multidimensional scaling method that aims to preserve local distances in the high-dimensional feature space. Both the distances within the subjects’ own data segment in the latent space, and the relative distances of within-subject distance to the average distance to others, referred to as (relative) self-distance score, were examined. The latter is defined as

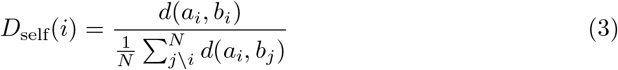

where *a*_*i*_ is the first data segment of subject *i* and *b*_*i*_ is the second data segment of subject i. The distance measure *d*(·) represents L1 distance. The nominator *d*(*a*_*i*_, *b*_*i*_) equals the absolute within-subject distance, and the denominator represents the average distance to others. A lower self-distance score means better within-subject stability. Conversely, when the absolute within-subject distance grows relative to the distance to others, the self-distance measure increases, indicating worse stability. The sex effect for within-subject distances and self-distance scores were tested using a standard t-test. To test the effect of the models’ input data to self-distance measure and average between subject distance, we fit Welch’s ANOVA. Post-hoc analysis was carried out using a nonparametric Games-Howell test.

To test the relationship between age and self-distance scores in the latent space, linear regression models were used. These were fit separately for females and males. We also tried quadratic regression models for the same data, but these did not yield significant relationships. Mantell’s permutation test [39] was employed to assess the Pearson correlation of distance matrices calculated from latent space projections produced by models trained in different sleep stages. The number of permutations to determine the significance was set to 999. We were especially interested in the stability of the latent space across sleep stage models, and its age dependency.

## Data availability

As the Finnish data protection legislation prohibits sharing the personal data of the participants, the EEG recordings are not available. The source code used in analysis of this work is freely available and hosted on GitHub (https://github.com/vernaverna/FABEEG).

## Acknowledgments

We thank the clinical neurophysiology technicians for the recording and marking of the EEG.

## Additional information

### Author contributions

V.H, M.L. and H.R. conceived and designed research, V.H. analyzed the data and drafted the manuscript; S.M. developed software; V.H., M.L. and H.R. interpreted the results and edited & revised the manuscript; L.L. and S.V. provided and curated the data and revised the manuscript; R.S. and S.M. revised the manuscript.

## Supplementary information

### Figures

**Figure 5 –figure supplement 1.**
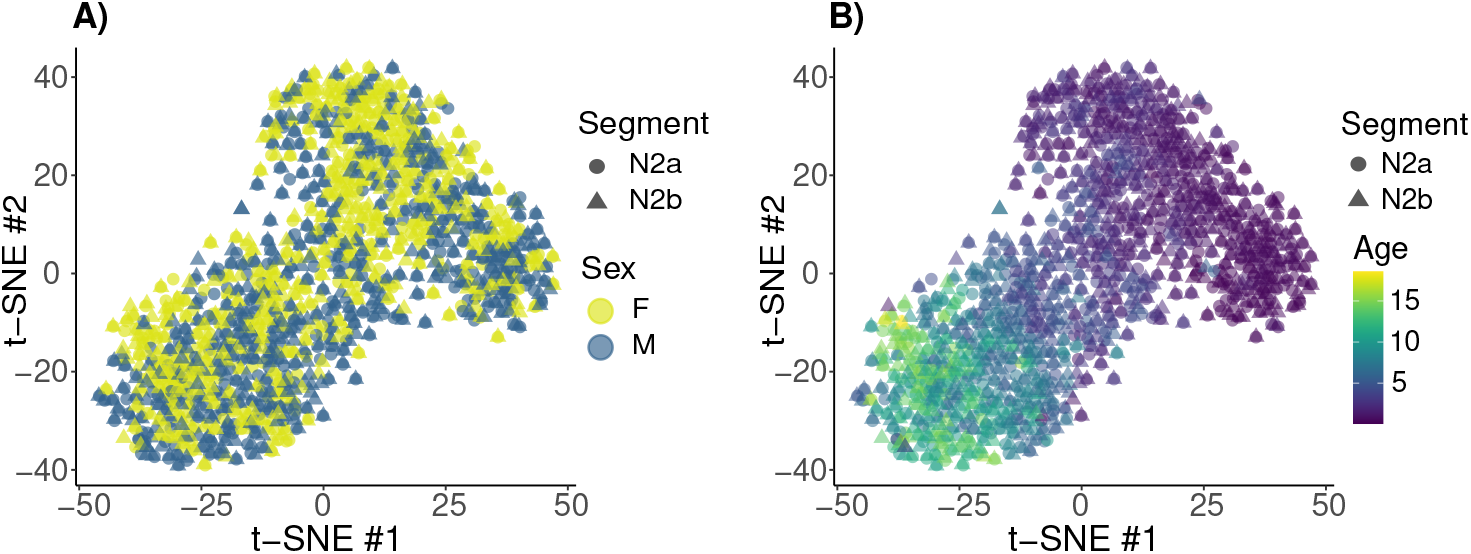
Two-dimensional t-SNE projection of the subjects projected in 30-dimensional latent mapping provided by BRRR (trained with N2 data). The first N2 segment is marked with a circle, the second with a triangle. As in Figure 5, we observe an age effect but not sex effect in the latent space.

**Figure 8 –figure supplement 1.**
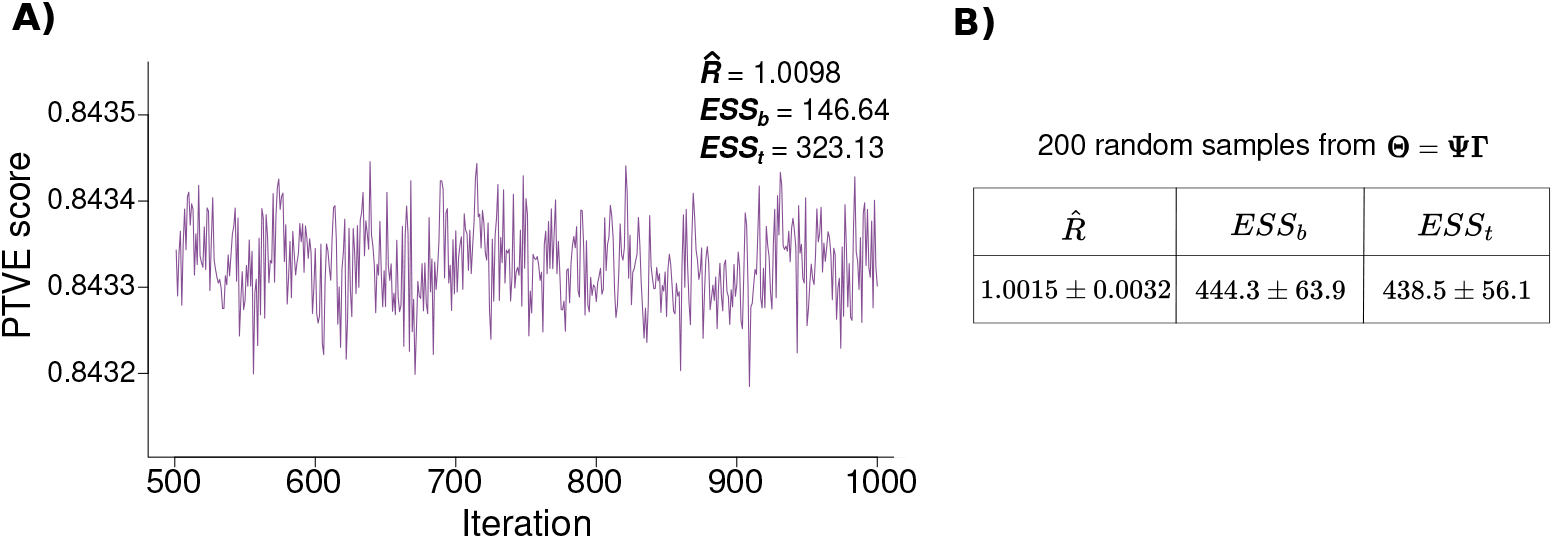
Model convergence diagnostics. Figure **A** depicts a trace plot of the model PTVE across iterations (after discarding the first half of the samples as burn-in period) along with Markov chain convergence diagnostics, demonstrating good convergence. Table **B** summarizes the converge diagnostics (mean ± SD) of 200 randomly sampled indices from the low-rank regression coefficient matrix.

## References

1. C. C. Aggarwal, A. Hinneburg, and D. A. Keim. On the surprising behavior of distance metrics in high dimensional space. In Database Theory—ICDT 2001: 8th International Conference London, UK, January 4–6, 2001 Proceedings 8, pages 420–434. Springer, 2001.

2. E. Amico and J. Goñi. The quest for identifiability in human functional connectomes. Scientific reports, 8(1):8254, 2018.

3. S. Berto, G.-Z. Wang, J. Germi, B. C. Lega, and G. Konopka. Human Genomic Signatures of Brain Oscillations During Memory Encoding. Cerebral Cortex, 28(5):1733–1748, 04 2017.

4. C. Binnie and P. Prior. Electroencephalography. Journal of Neurology, Neurosurgery & Psychiatry, 57(11):1308–1319, 1994.

5. M. Boersma, D. J. Smit, H. M. de Bie, G. C. M. Van Baal, D. I. Boomsma, E. J. de Geus, H. A. Delemarre-van de Waal, and C. J. Stam. Network analysis of resting state eeg in the developing young brain: structure comes with maturation. Human brain mapping, 32(3):413–425, 2011.

6. J. Buckelmüller, H.-P. Landolt, H. Stassen, and P. Achermann. Trait-like individual differences in the human sleep electroencephalogram. Neuroscience, 138(1):351–356, 2006.

7. F. T. Candelaria-Cook, I. Solis, M. E. Schendel, Y.-P. Wang, T. W. Wilson, V. D. Calhoun, and J. M. Stephen. Developmental trajectory of MEG resting-state oscillatory activity in children and adolescents: a longitudinal reliability study. Cerebral Cortex, 02 2022. bhac023.

8. C. J. Chu, J. Leahy, J. Pathmanathan, M. Kramer, and S. S. Cash. The maturation of cortical sleep rhythms and networks over early development. Clinical Neurophysiology, 125(7):1360–1370, 2014.

9. S. R. Cooper, J. J. Jackson, D. M. Barch, and T. S. Braver. Neuroimaging of individual differences: A latent variable modeling perspective. Neuroscience & Biobehavioral Reviews, 98:29–46, 2019.

10. L. Cragg, N. Kovacevic, A. R. McIntosh, C. Poulsen, K. Martinu, G. Leonard, and T. Paus. Maturation of eeg power spectra in early adolescence: a longitudinal study. Developmental science, 14(5):935–943, 2011.

11. J. da Silva Castanheira, H. D. Orozco Perez, B. Misic, and S. Baillet. Brief segments of neurophysiological activity enable individual differentiation. Nature communications, 12(1):5713, 2021.

12. J. da Silva Castanheira, A. I. Wiesman, J. Y. Hansen, B. Misic, S. Baillet, J. Breitner, J. Poirier, P. Bellec, V. Bohbot, M. Chakravarty, et al. The neurophysiological brain-fingerprint of parkinson’s disease. eBioMedicine, 105:105201, 2024.

13. L. De Gennaro, M. Ferrara, F. Vecchio, G. Curcio, and M. Bertini. An electroencephalographic fingerprint of human sleep. NeuroImage, 26(1):114–122, 2005.

14. M. Del Pozo-Banos, J. B. Alonso, J. R. Ticay-Rivas, and C. M. Travieso. Electroencephalogram subject identification: A review. Expert Systems with Applications, 41(15):6537–6554, 2014.

15. M. Demuru and M. Fraschini. Eeg fingerprinting: Subject-specific signature based on the aperiodic component of power spectrum. Computers in Biology and Medicine, 120:103748, 2020.

16. T. G. Dietterich. Approximate statistical tests for comparing supervised classification learning algorithms. Neural computation, 10(7):1895–1923, 1998.

17. T. Donoghue, M. Haller, E. J. Peterson, P. Varma, P. Sebastian, R. Gao, T. Noto, H. Lara, J. D. Wallis, R. T. Knight, et al. Parameterizing neural power spectra into periodic and aperiodic components. Nature neuroscience, 23(12):1655–1665, 2020.

18. T. Donoghue, N. Schaworonkow, and B. Voytek. Methodological considerations for studying neural oscillations. European journal of neuroscience, 55(11-12):3502–3527, 2022.

19. M. Eisermann, A. Kaminska, M.-L. Moutard, C. Soufflet, and P. Plouin. Normal eeg in childhood: from neonates to adolescents. Neurophysiologie Clinique/Clinical Neurophysiology, 43(1):35–65, 2013.

20. D. A. Engemann, O. Kozynets, D. Sabbagh, G. Lemâitre, G. Varoquaux, F. Liem, and A. Gramfort. Combining magnetoencephalography with magnetic resonance imaging enhances learning of surrogate-biomarkers. Elife, 9:e54055, 2020.

21. E. S. Finn, X. Shen, D. Scheinost, M. D. Rosenberg, J. Huang, M. M. Chun, X. Papademetris, and R. T. Constable. Functional connectome fingerprinting: identifying individuals using patterns of brain connectivity. Nature neuroscience, 18(11):1664–1671, 2015.

22. M. Fraschini, S. M. Pani, L. Didaci, and G. L. Marcialis. Robustness of functional connectivity metrics for eeg-based personal identification over task-induced intra-class and inter-class variations. Pattern Recognition Letters, 125:49–54, 2019.

23. A. Gelman, J. B. Carlin, H. S. Stern, D. B. Dunson, A. Vehtari, and D. B. Rubin. Bayesian data analysis. 3, 2013.

24. J. Gillberg, P. Marttinen, M. Pirinen, A. J. Kangas, P. Soininen, M. Ali, A. S. Havulinna, M.-R. Järvelin, M. Ala-Korpela, S. Kaski, et al. Multiple output regression with latent noise. Journal of Machine Learning Research, 2016.

25. A. Gramfort, M. Luessi, E. Larson, D. A. Engemann, D. Strohmeier, C. Brodbeck, R. Goj, M. Jas, T. Brooks, L. Parkkonen, et al. Meg and eeg data analysis with mne-python. Frontiers in neuroscience, page 267, 2013.

26. R. Guerrini. Epilepsy in children. The Lancet, 367(9509):499–524, 2006.

27. J. Haakana, S. Merz, S. Kaski, H. Renvall, and R. Salmelin. Bayesian reduced rank regression models generalizable neural fingerprints that differentiate between individuals in magnetoencephalography data. European Journal of Neuroscience, 59(9):2320–2335, 2024.

28. S. Haegens, H. Cousijn, G. Wallis, P. J. Harrison, and A. C. Nobre. Inter-and intra-individual variability in alpha peak frequency. Neuroimage, 92:46–55, 2014.

29. C. Hatlestad-Hall, R. Bruña, M. Liljeström, H. Renvall, K. Heuser, E. Taubøll, F. Maestú, and I. H. Haraldsen. Reliable evaluation of functional connectivity and graph theory measures in source-level eeg: How many electrodes are enough? Clinical Neurophysiology, 150:1–16, 2023.

30. A. Hyvarinen. Fast ica for noisy data using gaussian moments. In 1999 IEEE international symposium on circuits and systems (ISCAS), volume 5, pages 57–61. IEEE, 1999.

31. A.-K. Joechner, M. A. Hahn, G. Gruber, K. Hoedlmoser, and M. Werkle-Bergner. Sleep spindle maturity promotes slow oscillation-spindle coupling across child and adolescent development. Elife, 12:e83565, 2023.

32. K. A. Jones, B. Porjesz, L. Almasy, L. Bierut, A. Goate, J. C. Wang, D. M. Dick, Hinrichs, J. Kwon, J. P. Rice, J. Rohrbaugh, H. Stock, W. Wu, L. O. Bauer, D. B. Chorlian, R. R. Crowe, H. J. Edenberg, T. Foroud, V. Hesselbrock, S. Kuperman, J. Nurnberger Jr, S. J. O’Connor, M. A. Schuckit, A. T. Stimus, J. A. Tischfield, T. Reich, and H. Begleiter. Linkage and linkage disequilibrium of evoked eeg oscillations with chrm2 receptor gene polymorphisms: implications for human brain dynamics and cognition. International Journal of Psychophysiology, 53(2):75–90, 2004.

33. W. Klimesch. Eeg alpha and theta oscillations reflect cognitive and memory performance: a review and analysis. Brain research reviews, 29(2-3):169–195, 1999.

34. T. Koike-Akino, R. Mahajan, T. K. Marks, Y. Wang, S. Watanabe, O. Tuzel, and P. Orlik. High-accuracy user identification using eeg biometrics. In 2016 38th annual international conference of the IEEE engineering in medicine and biology society (EMBC), pages 854–858. IEEE, 2016.

35. D. La Rocca, P. Campisi, B. Vegso, P. Cserti, G. Kozmann, F. Babiloni, and F. D. V. Fallani. Human brain distinctiveness based on eeg spectral coherence connectivity. IEEE transactions on Biomedical Engineering, 61(9):2406–2412, 2014.

36. E. Leppäaho, H. Renvall, E. Salmela, J. Kere, R. Salmelin, and S. Kaski. Discovering heritable modes of meg spectral power. Human brain mapping, 40(5):1391–1402, 2019.

37. A. R. Levin, A. J. Naples, A. W. Scheffler, S. J. Webb, F. Shic, C. A. Sugar, M. Murias, R. A. Bernier, K. Chawarska, G. Dawson, et al. Day-to-day test-retest reliability of eeg profiles in children with autism spectrum disorder and typical development. Frontiers in integrative neuroscience, 14:21, 2020.

38. K. Li, J. Wang, S. Li, H. Yu, L. Zhu, J. Liu, and L. Wu. Feature extraction and identification of alzheimer’s disease based on latent factor of multi-channel eeg. IEEE Transactions on Neural Systems and Rehabilitation Engineering, 29:1557–1567, 2021.

39. N. Mantel. The detection of disease clustering and a generalized regression approach. Cancer research, 27(2 Part 1):209–220, 1967.

40. E. Maris and R. Oostenveld. Nonparametric statistical testing of eeg-and meg-data. Journal of neuroscience methods, 164(1):177–190, 2007.

41. A. F. Marquand, I. Rezek, J. Buitelaar, and C. F. Beckmann. Understanding heterogeneity in clinical cohorts using normative models: beyond case-control studies. Biological psychiatry, 80(7):552–561, 2016.

42. P. Marttinen, M. Pirinen, A.-P. Sarin, J. Gillberg, J. Kettunen, I. Surakka, A. J. Kangas, P. Soininen, P. O’Reilly, M. Kaakinen, et al. Assessing multivariate gene-metabolome associations with rare variants using bayesian reduced rank regression. Bioinformatics, 30(14):2026–2034, 2014.

43. A. Merkin, S. Sghirripa, L. Graetz, A. E. Smith, B. Hordacre, R. Harris, J. Pitcher, J. Semmler, N. C. Rogasch, and M. Goldsworthy. Do age-related differences in aperiodic neural activity explain differences in resting eeg alpha? Neurobiology of Aging, 121:78–87, 2023.

44. K. A. M. Pauls, P. Nurmi, H. Ala-Salomäki, H. Renvall, J. Kujala, and M. Liljeström. Human sensorimotor resting state beta events and aperiodic activity show good test–retest reliability. Clinical Neurophysiology, 163:244–254, 2024.

45. S. Perone, J. Palanisamy, and S. M. Carlson. Age-related change in brain rhythms from early to middle childhood: Links to executive function. Developmental science, 21(6):e12691, 2018.

46. S. Purcell, D. Manoach, C. Demanuele, B. Cade, S. Mariani, R. Cox, G. Panagiotaropoulou, R. Saxena, J. Pan, J. Smoller, et al. Characterizing sleep spindles in 11,630 individuals from the national sleep research resource. Nature communications, 8(1):1–16, 2017.

47. R Core Team. R: A Language and Environment for Statistical Computing. R Foundation for Statistical Computing, Vienna, Austria, 2020.

48. M. Sankupellay, S. Wilson, H. Heussler, C. Parsley, M. Yuill, and C. Dakin. Characteristics of sleep eeg power spectra in healthy infants in the first two years of life. Clinical neurophysiology, 122(2):236–243, 2011.

49. E. Sareen, S. Zahar, D. Van De Ville, A. Gupta, A. Griffa, and E. Amico. Exploring meg brain fingerprints: Evaluation, pitfalls, and interpretations. NeuroImage, 240:118331, 2021.

50. N. Schaworonkow and B. Voytek. Longitudinal changes in aperiodic and periodic activity in electrophysiological recordings in the first seven months of life. Developmental cognitive neuroscience, 47:100895, 2021.

51. S. Shahab, B. H. Mulsant, M. L. Levesque, N. Calarco, A. Nazeri, A. L. Wheeler, G. Foussias, T. K. Rajji, and A. N. Voineskos. Brain structure, cognition, and brain age in schizophrenia, bipolar disorder, and healthy controls. Neuropsychopharmacology, 44(5):898–906, 2019.

52. S. J. Smith. Eeg in the diagnosis, classification, and management of patients with epilepsy. Journal of Neurology, Neurosurgery & Psychiatry, 76(Suppl 2):ii2–ii7, 2005.

53. P. Sorrentino, R. Rucco, A. Lardone, M. Liparoti, E. T. Lopez, C. Cavaliere, Soricelli, V. Jirsa, G. Sorrentino, and E. Amico. Clinical connectome fingerprints of cognitive decline. NeuroImage, 238:118253, 2021.

54. K. T. Tapani, P. Nevalainen, S. Vanhatalo, and N. J. Stevenson. Validating an svm-based neonatal seizure detection algorithm for generalizability, non-inferiority and clinical efficacy. Computers in Biology and Medicine, 145:105399, 2022.

55. L. Tarokh, M. A. Carskadon, and P. Achermann. Trait-like characteristics of the sleep eeg across adolescent development. Journal of Neuroscience, 31(17):6371–6378, 2011.

56. P. M. Thompson, T. Ge, D. C. Glahn, N. Jahanshad, and T. E. Nichols. Genetics of the connectome. Neuroimage, 80:475–488, 2013.

57. M. Tröndle, T. Popov, S. Dziemian, and N. Langer. Decomposing the role of alpha oscillations during brain maturation. ELife, 11:e77571, 2022.

58. S. A. Valizadeh, F. Liem, S. Mérillat, J. Hänggi, and L. Jäncke. Identification of individual subjects on the basis of their brain anatomical features. Scientific reports, 8(1):5611, 2018.

59. D. Van De Ville, Y. Farouj, M. G. Preti, R. Liégeois, and E. Amico. When makes you unique: temporality of the human brain fingerprint. Science advances, 7(42):eabj0751, 2021.

60. L. Van der Maaten and G. Hinton. Visualizing data using t-sne. Journal of machine learning research, 9(11), 2008.

61. A. Vehtari, A. Gelman, D. Simpson, B. Carpenter, and P.-C. Bürkner. Rank-normalization, folding, and localization: An improved r for assessing convergence of mcmc (with discussion). Bayesian analysis, 16(2):667–718, 2021.

62. D. Vidaurre, A. J. Quinn, A. P. Baker, D. Dupret, A. Tejero-Cantero, and M. W. Woolrich. Spectrally resolved fast transient brain states in electrophysiological data. Neuroimage, 126:81–95, 2016.

63. T. J. Whitford, C. J. Rennie, S. M. Grieve, C. R. Clark, E. Gordon, and L. M. Williams. Brain maturation in adolescence: concurrent changes in neuroanatomy and neurophysiology. Human brain mapping, 28(3):228–237, 2007.

64. A. I. Wiesman, J. da Silva Castanheira, and S. Baillet. Stability of spectral estimates in resting-state magnetoencephalography: Recommendations for minimal data duration with neuroanatomical specificity. Neuroimage, 247:118823, 2022.

65. S. Zhang, W. Yang, H. Mou, Z. Pei, F. Li, and X. Wu. An overview of brain fingerprint identification based on various neuroimaging technologies. IEEE Transactions on Cognitive and Developmental Systems, 2023.

66. H. Zhu, Z. Khondker, Z. Lu, and J. G. Ibrahim. Bayesian generalized low rank regression models for neuroimaging phenotypes and genetic markers. Journal of the American Statistical Association, 109(507):977–990, 2014.

